# Phage defense and genome editing using novel retrons sourced from isolated environmental bacteria

**DOI:** 10.1101/2025.01.29.635429

**Authors:** Kazuo L Nakamura, Karen Zhang, Mario R Mestre, Matías Rojas-Montero, Seth L Shipman

**Affiliations:** Gladstone Institute of Data Science and Biotechnology, San Francisco, CA, USA; Graduate Program in Bioengineering, University of California, San Francisco and Berkeley, CA, USA; Section of Microbiology, Department of Biology, University of Copenhagen, Copenhagen, Denmark; Department of Bioengineering and Therapeutic Sciences, University of California, San Francisco, CA, USA; Chan Zuckerberg Biohub, San Francisco, CA, USA

## Abstract

Retrons are bacterial immune systems that protect a bacterial population against phages by killing infected hosts. Retrons typically comprise a reverse transcriptase, a template noncoding RNA that is partially reverse transcribed into RT-DNA, and a toxic effector. The reverse transcriptase, noncoding RNA, and RT-DNA complex sequester the toxic effector until triggered by phage infection, at which point the toxin is released to induce cell death. Due to their ability to produce single-stranded DNA in vivo, retrons have also been engineered to produce donor templates for genome editing in both prokaryotes and eukaryotes. However, the current repertoire of experimentally characterized retrons is limited, with most retrons sourced from clinical and laboratory strains of bacteria. To better understand retron biology and natural diversity, and to expand the current toolbox of retron-based genome editors, we developed a pipeline to isolate retrons and their bacterial hosts from a variety of environmental samples. Here, we present six of these novel retrons, each isolated from a different host bacterium. We characterize the full operon of these retrons and test their ability to defend against a panel of *E. coli* phages. For two of these retrons, we further unravel their mechanism of defense by identifying the phage genes responsible for triggering abortive infection. Finally, we engineer these retrons for genome editing in *E. coli*, demonstrating their potential use in a biotechnological application.

## INTRODUCTION

Retrons are multi-component bacterial immune systems that trigger abortive infection in response to phage. The components of these systems include a reverse transcriptase (RT), a noncoding RNA (ncRNA) that is partially reverse transcribed to a hybrid RNA/DNA molecule, and one or more effector proteins that typically function as toxins to the retron host^1-3^. The toxicity of the effector protein(s) is neutralized in the presence of the RT and reverse-transcribed DNA (RT-DNA), but perturbation of the RT-DNA by phage-encoded elements can release this neutralization, leading to abortive infection. Beyond their natural function in phage defense, retrons have also be repurposed into components of biotechnology^4-6^. For instance, retron ncRNA can be modified so that reverse transcription produces template DNA for genome editing^7-18^.

Retrons were initially discovered in Myxobacteria when researchers noticed unexplained bands running on acrylamide gels of ∼120 nucleotides in length from total nucleic acid preparations^19^. Subsequent studies determined that these bands were single-stranded DNA (ssDNA), produced by a retron RT from a retron ncRNA^20^. More recently, thousands of retrons have been bioinformatically predicted in diverse bacterial species and categorized into clades and subtypes^21^. These retrons have different effector proteins and mechanisms, as well as widely varying RT-DNA production and utility in gene editing^22^.

Despite this, only a handful of retrons have been studied in the context of their natural host, and retron defense against phages has only been studied in *E. coli*^*3,23-25*^, *B. subtilis*^*26*^, and *S. enterica*^*1*^. Furthermore, prior studies have been largely limited to retrons from clinical and laboratory strains of bacteria with few environmental examples (e.g. *Vibrio mimicus*^*27*^).

In this study, we develop an approach to identify novel retrons from bacteria isolated from soil and water samples by screening for RT-DNA bands using DNA isolation and polyacrylamide gel electrophoresis. Here, we add six new environmental retron hosts. We demonstrate that four of six naturally sourced retrons can be ported into *E. coli* to produce RT-DNA and two exhibit defense against an *E. coli* phage, Bas51. By sequencing phage escapees, we determine that the Bas51 N6 adenine methyltransferase is a trigger for these two retrons. We also demonstrate that these environmentally sourced retrons can be used for genome editing in *E. coli*. This process for isolating and identifying novel retrons can be used to expand our understanding of retron biology by providing more natural hosts for study and aid in the development of new tools for genome editing.

## RESULTS

### Isolation of Retron-Bearing Hosts from Environmental Samples

To isolate new retron hosts from the environment, soil and water samples were collected from a range of climates and environments (**Supplemental Table 1**). Soil samples were added to LB media and broken down by vortexing with beads. Rocks and soil debris were removed by centrifugation through a filter. The resulting media containing soil microbes was plated on LB agar. Colonies were picked and grown for 16h in LB media, then passaged at a 1:100 dilution and grown for an additional 7h in LB media (**Fig 1a**). Water samples were plated directly and colonies were subcultured using the steps described above. All samples in this study were initially cultured at 37° C, but the *S. quinivorans* and *H. chinensis* samples were cultured at 30° C in later experiments after exhibiting faster growth at this temperature.

**Figure 1.**
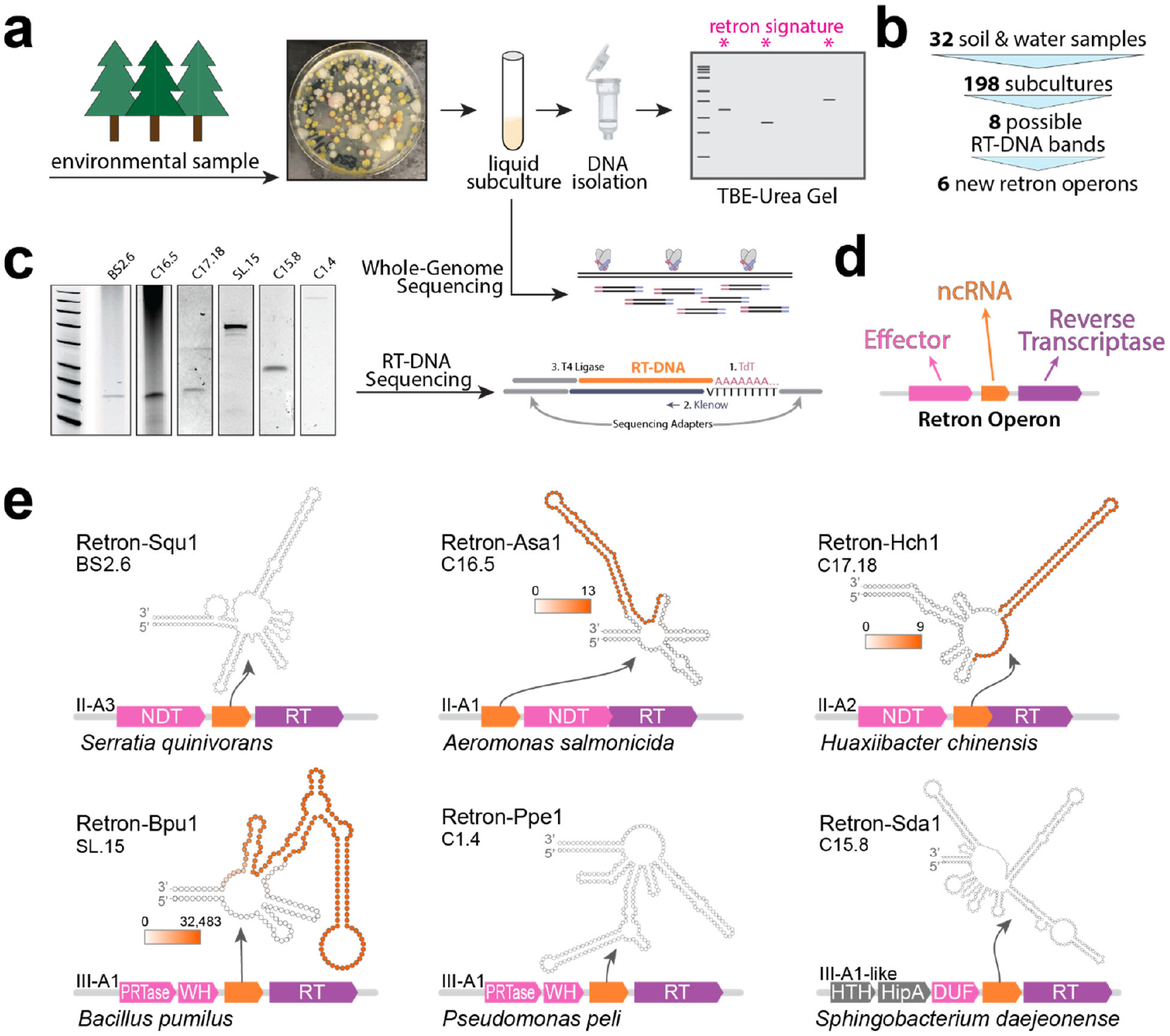
Isolation of Retron-Bearing Hosts from Environmental Samples. **A)** Schematic of isolation of bacteria from soil and water samples and identification of possible retron-bearing hosts using PAGE analysis. **B)** Overview of sampling, screening of colonies, and identification of retron hosts. **C)** Retron RT-DNA bands from PAGE analyses (uncropped gels included as Supplementary Figure 1). **D)** Schematic of canonical retron operon. **E)** Six new retron operons, identifying the retron subtype, host species, and predicted two-dimensional structure of the ncRNA. For retrons where RT-DNA sequencing is available, sequencing coverage is plotted onto ncRNA in an orange heatmap (heatmap legend indicated number of sequencing reads mapped).

We identified retron-bearing hosts by the presence of the characteristic retron ssDNA that runs at 40-180 nucleotides on a polyacrylamide gel. To do so, we prepped total DNA from the 7h cultures using Qiagen miniprep kits and ran this DNA on TBE-Urea gels.

In total, 13 water samples and 19 soil samples were collected. DNA was isolated from 198 subcultures and run on gels. 29 of the subcultures exhibited possible bands on initial gels. Seven subcultures were not followed up because they duplicated band positions from other bacteria originating from the same environmental sample. The remaining 22 subcultures with possible bands were recultured and rechecked. Eight of these subcultures could not be recovered from glycerol stocks and six subcultures showed no distinct bands upon rechecking, leaving eight subcultures with reliable RT-DNA production, all originating from different environmental samples. We sequenced genomes of the remaining eight subcultures. In six of the sequenced genomes, we found putative retron operons (**Fig 1b, Supplemental Table 3**).

Bacterial subcultures showing a putative retron RT-DNA band underwent both whole genome sequencing, as well as single stranded DNA isolation followed by RT-DNA sequencing (**Fig 1c**). Whole genome data was analyzed using the Prokaryotic Antiviral Defense Locator (PADLOC)^28^ tool, which identifies retron operons and the genes encoding the effector protein, noncoding RNA, and reverse transcriptase, which are present in most retron subtypes (**Fig 1d**). We also analyzed whole genome contigs using PubMLST, which called the species of each subculture based on genes encoding ribosome protein subunits. In total, PADLOC predicted six novel retron operons from the II-A and III-A subtypes in the *Serratia quinivorans* (Retron-Squ1), *Aeromonas salmonicida* (Retron-Asa1), *Huaxiibacter chinensis* (Retron-Hch1), *Bacillus pumilus* (Retron-Bpu1), *Pseudomonas peli* (Retron-Ppe1), and *Sphingobacterium daejeonense* (Retron-Sda1) genomes, each of which is the first experimentally studied retron in these species (**Fig 1e**). The noncoding RNA sequences of Retrons-Squ1, -Asa1, -Hch1, -Sda1, and -Ppe1 were predicted by PADLOC. In the case of Retron-Bpu1, the noncoding RNA was refined after RT-DNA sequencing data mapped beyond the noncoding RNA that PADLOC predicted. In three cases, we were able to sequence the retron RT-DNA and plot coverage onto the ncRNA (coverage in orange).

### Retron-Sda1 is an example of a previously unknown retron type

In the case of Retron-Sda1, the identified ncRNA was classified as a type III-A1 ncRNA. Surprisingly, instead of being co-located with the PRTase and DNA-binding domain-containing proteins that characterize retrons type III, Retron-Sda1 was co-located with two genes encoding proteins for HipA-like kinase and DUF3037 domains, respectively (**Fig 1e** and **Fig 2c,e**). Given the lack of the usual Type III effectors, we decided to investigate this case in more depth. For this, we queried the NCBI nr database in search of homologs for both the RT and the DUF3037 proteins, performed multiple sequence alignments and phylogenetic analyses, and retrieved the genomic neighborhoods using the NCBI Entrez API and annotated the defense systems using PADLOC. Interestingly, DUF3037 was found to be strongly associated with HipA-like containing proteins (Supplementary File 1). This two-gene system was identified as a putative defense system PDC-M66 (**Fig 2a**) across numerous homologs of the tree. A similar domain architecture was found recently in the experimentally-validated system DS-6^29^. In the RT tree, the association of the RT with HipA/DUF3037 occurred at least in 6 different cases (**Fig 2b,c** and **Supplementary File 2**) in two different clades (**Fig 2b**). One occurrence is found in species from the Bacteroidota phylum, and the other one is in species from the Pseudomonadota phylum (**Fig 2b,c**). Furthermore, we found that the C-terminal region of the DUF3037 shares sequence similarity to the DNA-binding protein from close Type III-A1 retrons (**Fig 2d**), likely representing a recent evolutionary event where the PRTase effector has been replaced by a predicted functional system (PDC-M66) formed by HipA and DUF3037. Although not very abundant, this case may represent a nascent association where the retron has captured new effectors, in an scenario similar to what occurred in Retron types II-A1 and I-A, where the NDT effector was replaced by a Septu defense system^21^.

**Figure 2.**
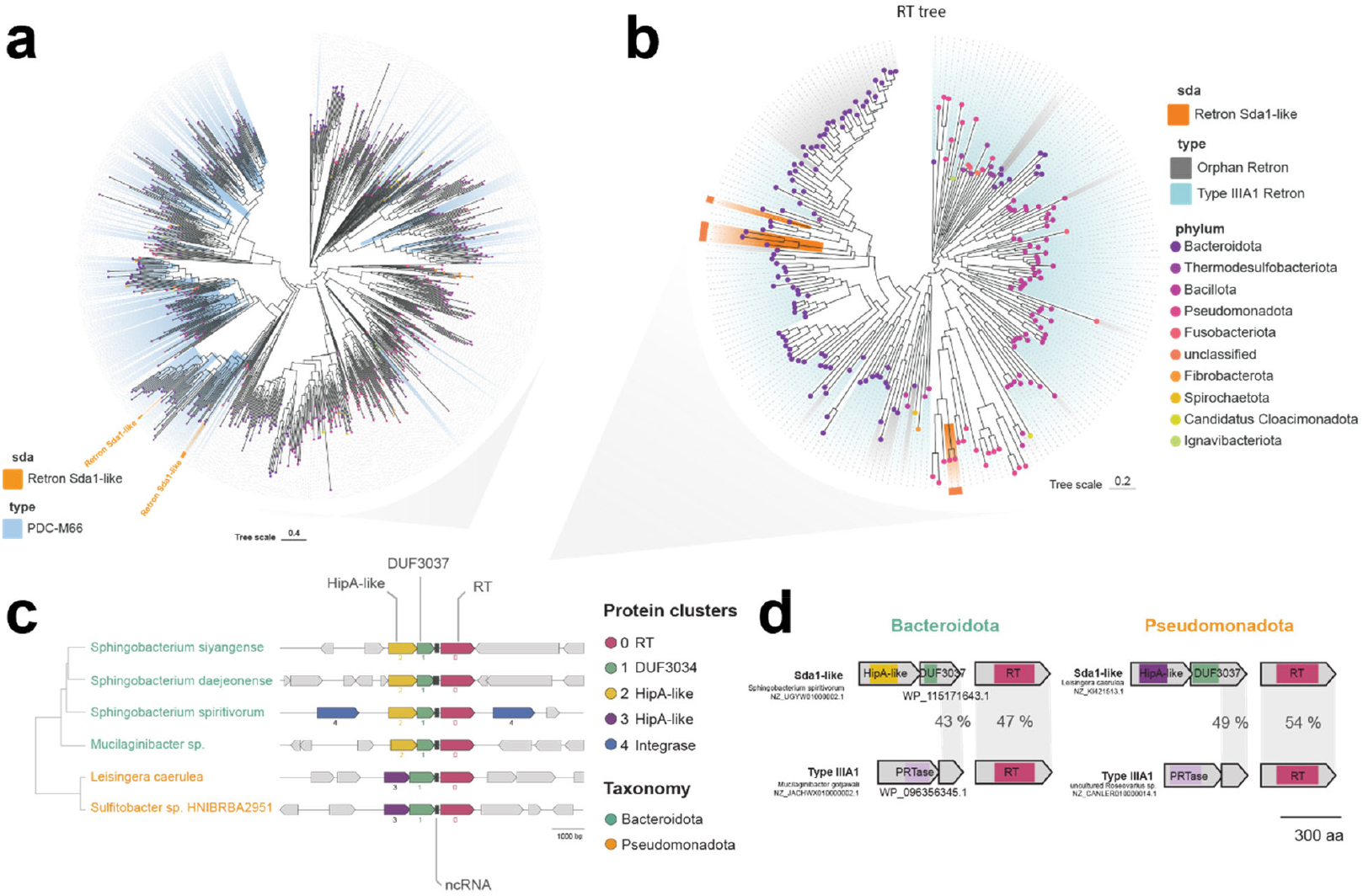
Retron-Sda1 is an example of a previously unknown retron type. **A)** Phylogenetic analysis of DUF3037-containing genomic neighborhoods. **B)** Phylogenetic analysis of RT homologs. **C)** Six examples of retron RT associated with DUF3034 and HipA-like proteins drawn from non-neighboring branches of the tree in b. **D)** Sequence homology between the c-terminus of the DUF3037 gene and the PRTase effector normally associated with Type III-AI retrons.

### Naturally Sourced Retrons Can Produce RT-DNA In E. Coli

To validate that the retron systems we identified were complete, we amplified them from the genomes of their natural host, cloned them onto plasmids, and transformed them for expression in *E. coli* bSLS114 (a derivative of BL21-AI with endogenous Retron-Eco1 removed) downstream of an inducible T7/lac promoter (**Fig 3a**). In cloning retron operons, we included 100-150 bases before the start codon of the first gene.

**Figure 3.**
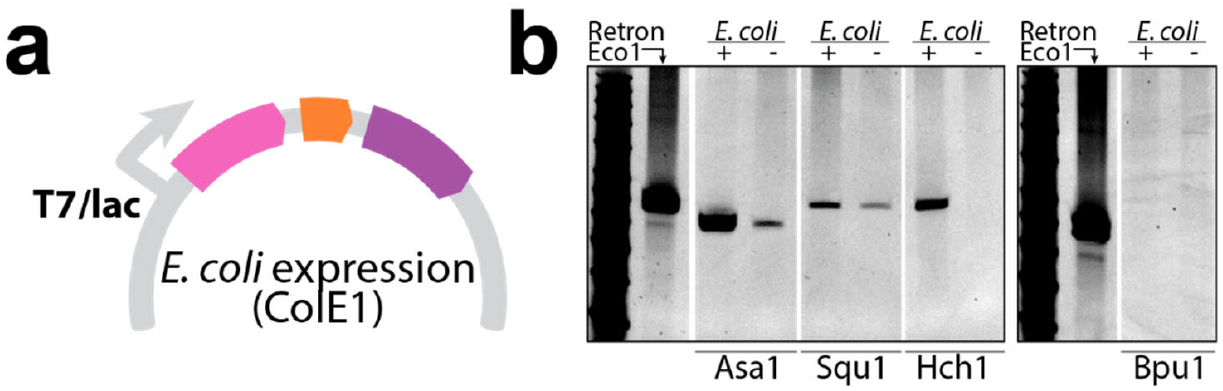
Naturally Sourced Retrons Can Produce RT-DNA In E. Coli. **A)** Schematic of plasmid for retron expression in E. coli. **B)** Images from PAGE analyses of RT-DNA production from naturally sourced retrons in induced and uninduced conditions (uncropped gels included as Supplementary Figure 2).

We analyzed RT-DNA production in both uninduced and induced conditions, to see if the systems were competent for production and if the systems carried their own constitutive promoters that are active in *E. coli*. Similar to analysis from within the natural host, cultures were grown at 37°C in LB media overnight, then passaged at a 1:100 dilution and grown for an additional 7 hours the following day. DNA was isolated using a Qiagen miniprep then analyzed on a TBE-Urea gel (**Fig 3b**).

Retron-Ppe1, identified in *Pseudomonas*, was not studied in *E. coli* because it could not be transformed, despite multiple attempts even in the presence of 1% glucose as a repressor of the inducible promoter. We presume that this retron is constitutively expressed and toxic to *E. coli*, perhaps due to the presence of a trigger for this retron in *E. coli*, or the absence of a neutralizing factor from the natural host. We also did not further study the III-A1-like Retron-Sda1.

Retron-Bpu1 exhibited RT-DNA production only in the uninduced sample but none when induced, suggesting there is a biological limitation affecting the retron’s RT-DNA production. However, all three type II-A retrons were successfully induced and exhibited stronger bands.

All four samples exhibited bands at the expected positions, which indicates that they were responsible for the initial RT-DNA bands in Fig 1a and that both the correct RT gene and msr-msd are included on the plasmids.

### Retrons Exhibit Defense Against Bas51 phage

To better understand how naturally sourced retrons function as defense systems, we tested their defense against T4, T5, lambda, and Bas51 phages in *E. coli*. While these retrons may defend against other phages in their natural hosts, retrons often provide defense against phages when ported into a new species as well.

Cultures of *E. coli* MG1655 transformed with Retrons-Asa1, -Squ1, -Hch1, and -Bpu1 (same plasmids as **Fig 3**) were plated in LB agar. Phages were then titrated and spotted, and plates were left at 37°C overnight for plaques to form.

None of the four naturally sourced retrons we tested exhibited defense against the T4 phage (**Fig 4a,b**). However, the Asa1 retron showed a subtle phenotype against T5, causing smaller plaques compared to the group with no retron (**Fig 4d**), although total plaque counts were comparable (**Fig 4c**) across all groups. Lambda phage plaque sizes with and without retrons and the quantity of plaques were similar across all groups. (**Fig 4e,f**). Asa1 exhibited a strong trend of defense against Bas51 at p = ∼0.6 and Squ1 exhibited significant defense against the same phage (**Fig 4g,h**). Retrons-Hch1 and -Bpu1 exhibited no defense against any of the tested phages.

**Figure 4.**
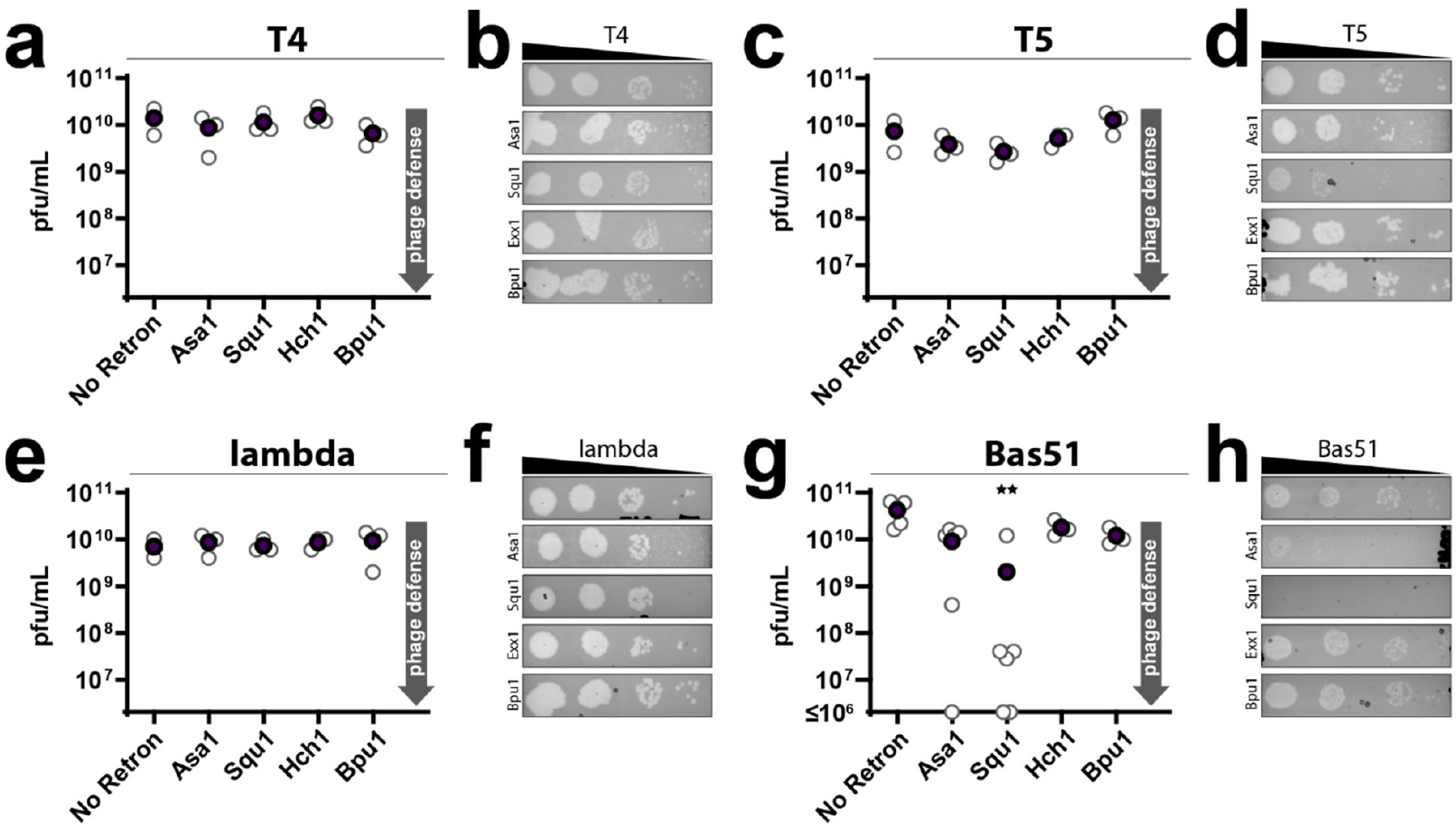
Retrons Exhibit Defense Against Bas51 phage. **A)** PFU/mL of T4 phage on strains containing constructs with naturally sourced retrons (Kruskal-Wallis test P=0.3385). **B)** Titration of T4 on strains containing constructs with naturally sourced retrons. **C)** PFU/mL of T5 phage on strains containing constructs with naturally sourced retrons (Kruskal-Wallis test P=0.0963). **D)** Titration of T5 on strains containing constructs with naturally sourced retrons. **E)** PFU/mL of lambda phage on strains containing constructs with naturally sourced retrons (Kruskal-Wallis test P=0.0984). **F)** Titration of lambda phage on strains containing constructs with naturally sourced retrons.. **G)** PFU/mL of Bas51 phage on strains containing constructs with naturally sourced retrons (Kruskal-Wallis test P=0.0047, follow-up Dunn’s test corrected for multiple comparisons: C16.5 vs no retron P=0.0631, BS2.6 vs no retron P=0.0009).

### Defense Mechanisms of Retron-Asa1 and -Squ1

We next identified the mechanism of Bas51 triggering mechanism for Retrons-Asa1 and -Squ1. Cells transformed with retrons as above were plated and Bas51 was spotted at low concentrations, allowing phages with mutations in genes triggering the retron to escape retron defense and form plaques.

We used nanopore sequencing to identify potential trigger genes in mutant phage escapees, sequencing five escapees of Retron-Asa1 and five of Retron-Squ1. When analyzing data, we ignored mutations in polynucleotide stretches (known to be enriched for nanopore errors) and synonymous mutations. All six distinct escapees that we sequenced contained mutations in an adenine methyltransferase gene (**Fig 5a**). Additionally, four of six contained premature stop mutations in a glycosidase gene (**Fig 5b**).

**Figure 5.**
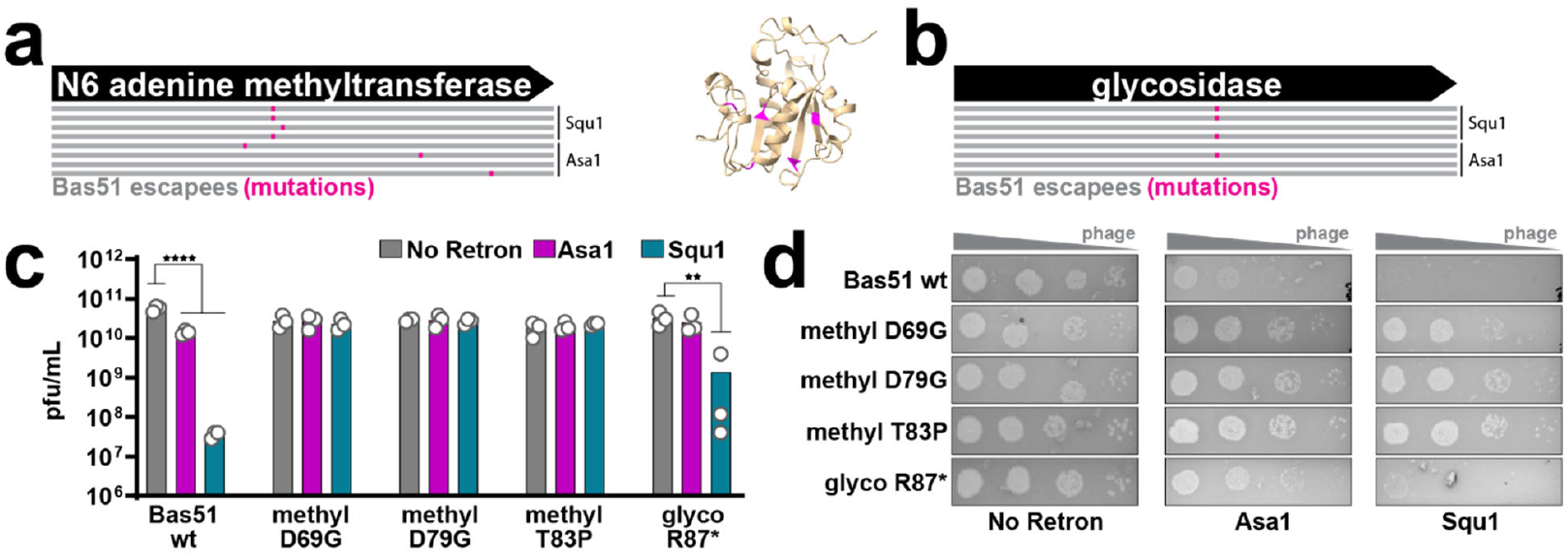
Defense Mechanisms of Retron-Asa1 and -Squ1. **A)** Left, phage escapee mutations in Bas 51 N6 adenine methyltransferase gene on strains containing Asa1 and Squ1 constructs; right, predicted 3D structure (AlphaFold3) of methyltransferase with mutations in pink. **B)** Phage escapee mutations in Bas 51 glycosidase gene on strains containing Asa1 and Squ1 constructs. **C)** PFU/mL of Bas51 phage with mutations in methyltransferase and glycosidase genes on strains containing Asa1 and Squ1 constructs (Two-way ANOVA effect of retron P<0.0001, phage P=0.1244, and interaction P<0.0001; follow-up testing with Tukey’s, corrected for multiple comparisons, wild-type Bas51 no retron versus Retron-Asa1 P<0.0001 and –Squ1 P<0.0001, glyco R87* no retron versus Squ1 P=0.0001, all other conditions ns. **D)** Titration of Bas51 phage with mutations in methyltransferase and glycosidase genes on strains containing Retron-Asa1 and -Squ1 constructs.

To determine whether either the methyltransferase or glycosidase gene functioned as a trigger, we initially tried expressing the genes individually in the presence or absence of the retrons to see if either resulted in a growth defect in the presence of the retron. However, we found that the methyltransferase was toxic to cells even in the absence of the retron.

To avoid the issue of methyltransferase toxicity, we instead introduced mutations in the methyltransferase and glycosidase genes in Bas51. By making these mutations precisely on a wild type phage background, we could disentangle potential effects of the methyltransferase and glycosidase (which were often both mutated in escapees) and remove the confounding influence of other mutations that were present in escapees. In separate phages, we precisely introduced three of the escapee mutations in the N6 adenine methyltransferase (D69G, D79G, and T83P) and the premature stop mutation in the glycosidase using a recombitron approach^30^.

Wild type Bas 51 propagates significantly less in both the Retron-Asa1 and -Squ1-expressing strains compared to a strain with no retron. However, mutating the methyltransferase gene at any of three tested positions eliminated defense by both Asa1 and Squ1, resulting in normal plaque formation (**Fig 4c,d**). This indicates that the Bas 51 N6 adenine methyltransferase gene triggers both retrons. Introducing a premature stop mutation in the glycosidase gene also eliminated the phage defense phenotype of Retron-Asa1, but not Retron-Squ1 (**Fig 5c,d**).

### Precise Editing Using Recombitrons Based On Naturally-Sourced Retrons

The recombitron approach that we used to edit Bas51 and identify its methyltransferase as the trigger is enabled by repurposing retron components into biotechnology for genome editing. In recombitrons, modified retron ncRNA is reverse transcribed by the retron RT to produce RT-DNA that serves as a recombineering donor, where the RT-DNA is incorporated into the lagging strand during DNA replication, precisely changing the genome in the process^7-9,15,31^. This approach increases editing efficiencies compared to producing template DNA *in vitro* then delivering it to a cell and is particularly useful when editing phage genomes because the recombineering donor is produced in high abundance within the host^30^. We have found that this technology can be built on different retrons, but that some retrons support more efficient editing than others^22^. To determine whether naturally sourced retrons could be used to design efficient editors, we engineered them to produce templates for recombineering of the *lacZ* gene in *E. coli* **(Fig 6a)**.

**Figure 6.**
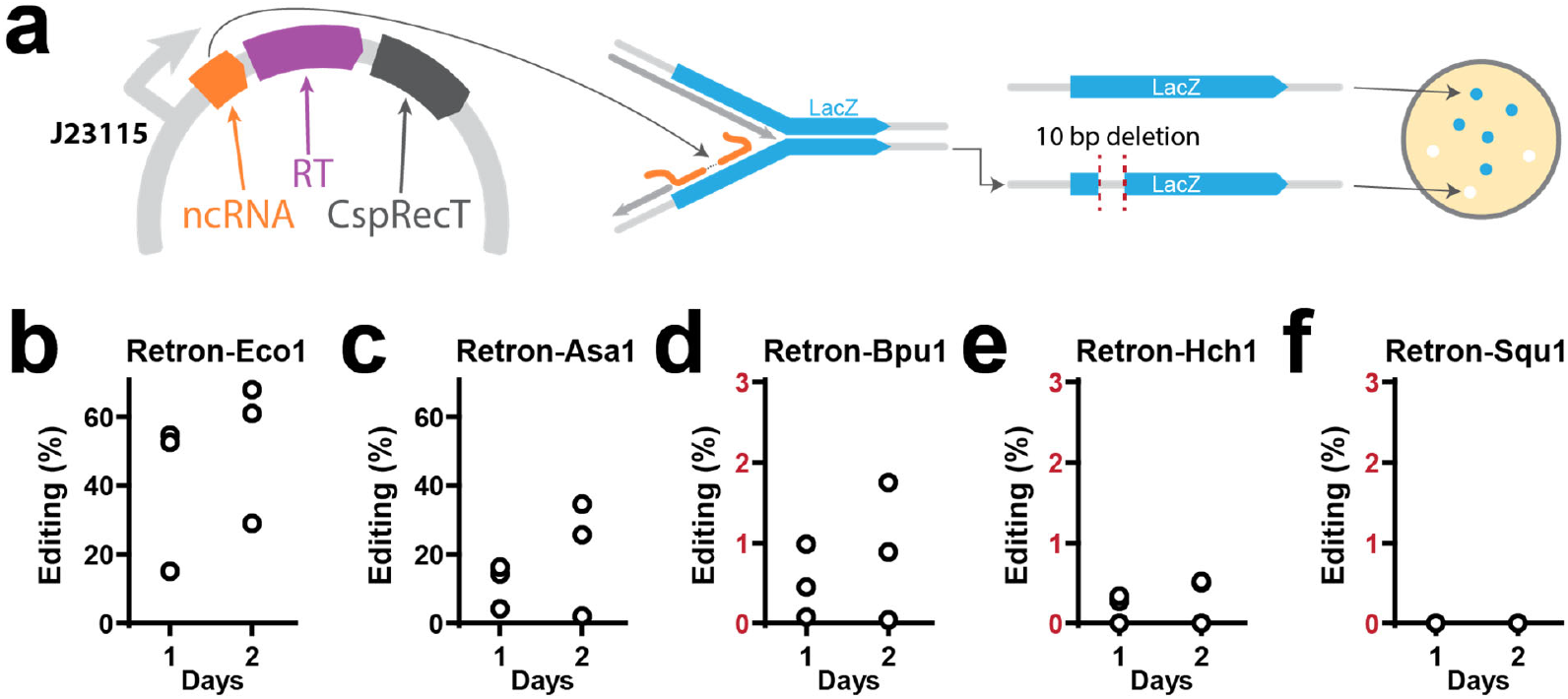
Precise Editing Using Recombitrons Based On Naturally-Sourced Retrons. **A**)Left, schematic of plasmid for recombineering in E. coli; right, schematic of recombineering LacZ gene with 10 bp deletion. B)Percent of cells with LacZ gene precisely edited using the Eco1 recombitron after culturing for 1 or 2 days, quantified using Illumina sequencing. **C)** Experiments and plots identical to **B**, but using the Asa1 (**C**), Bpu1 (**D**), Hch1 (**E**), and Squ1 (**F**) recombitrons.

Naturally sourced retron RT-DNAs were modified to encode a template for a 10 bp deletion in the *lacZ* gene. For retrons without RT-DNA data, the RT-DNA was assumed to cover the longest stem-loop in the noncoding RNA secondary structure. Retrons were expressed on a plasmid also containing a gene for CspRecT, a single stranded DNA annealing protein that enables efficient recombineering from the reverse transcribed donor. Based on prior studies, donor sequences were inserted roughly 10 base pairs out from the base of the main RT-DNA stem loop, replacing the remainder of the stem and the loop. Editing data was quantified by Illumina sequencing.

The Retron-Eco1 editor, which uses a well-studied retron from *E. coli* as a scaffold, exhibited higher editing rates than editors using naturally sourced retrons (**Fig 6b**). Among editors based on naturally sourced retrons, the Retron-Asa1 editor was the most efficient and made deletions in ∼30% of cells by the second culture round (**Fig 6c**). The remaining three retron editors exhibited much lower editing rates (below 2% after one day) (**Fig 6d,e,f**). Retron-Asa1 recombitron’s editing rates are likely partly a result of its RT-DNA production in E. coli, which far exceeds the other naturally sourced retrons.

## DISCUSSION

Though many retrons have been studied in *E. coli*, few have been identified from environmental samples or studied in their native bacterial hosts. This study describes a new approach to isolate retrons and their native hosts from environmental soil and water samples by screening for RT-DNA bands on polyacrylamide gels. This approach expands the collection of known retron hosts and could enable studies of retrons that cannot be transformed into *E. coli* or who function differently outside of their natural host bacterium.

Of the six novel retrons identified, five fit the II-A and III-A subtypes and one bears similarity to the III-A subtype but contains different accessory genes. While it is possible that not all retron subtypes produce RT-DNA constitutively, the majority do and should be identifiable when screening for RT-DNA bands. The overrepresentation of type II-A and III-A retrons may reflect selection due to culturing conditions favoring bacteria harboring these subtypes and the overall prevalence of these subtypes in nature.

PADLOC identified retron operons in whole genome sequencing data of most samples where clear, possible RT-DNA bands were visible on polyacrylamide gels. However, PADLOC found no retrons or retron RT genes in two of the nine sequenced genomes. These two genomes, which are identical, may contain retrons in regions where whole genome sequencing coverage was poor. It is also possible that an uncharacterized retron subtype could be present or another genetic element besides a retron may be responsible for the bands.

The discovery of Retron-Sda1 demonstrates that retron diversity extends beyond the thirteen currently recognized subtypes^21^. Interestingly, Retron-Sda1 contains a type III retron RT gene but two entirely different accessory genes. Where typical type III retron operons contain a phosphoribosyltransferase-like gene, Sda1 has a gene containing a HipA-like kinase domain and DUF3037 domain that have been predicted to function as a defense system on their own^29^. This new association likely represents a new retron type, in which the RT and the ncRNA may sense the phage signal, and the HipA/DUF3037 pair likely encodes effectors with toxic functions that are activated after sensing. The identification of Retron-Sda1 shows how the approach outlined in this study can capture novel retrons with unique features. In such cases, the presence of a retron RT gene may be a useful indicator for identifying retrons whose operons are not easily classified into the thirteen subtypes and would be challenging to identify using PADLOC and other bioinformatics tools.

We also show that the N6 adenine methyltransferase in Bas 51 acts as a trigger for Retron-Squ1 and both the N6 adenine methyltransferase and glycosidase contribute to triggering Retron-Asa1. This suggests that these naturally sourced retrons are triggered by methylation of the RT-DNA, as has been shown for other retrons^1^, whereas the glycosidase perturbs the retron effector bound complex via an unknown mechanism. The presence of two Bas 51 trigger genes required for activation of Retron-Asa1 may provide an additional safeguard against the retron being triggered in the absence of phage.

While most naturally sourced retrons isolated in this study are not efficient editors in *E. coli*, the Asa1 recombitron exhibits gene editing potential. Further research could explore the gene editing capabilities of recombitrons in other cell types or conditions more similar to their native host bacteria and environment.

## METHODS

### Isolating Bacteria from Soil and Water Samples

Small samples of soil (∼2.5 grams) and water (∼10 mL) were collected from glacial lakes and streams as well as alpine mountains, woodlands, and parks. Microbes were isolated from soil samples using the Zymo fecal/soil microbe kit, which removed dirt and rocks. Water and soil processed with the Zymo kit were then plated on plain LB and allowed to grow overnight or until colonies formed at 37°C.

### Identifying RT-DNA Bands on Polyacrylamide Gel

Colonies were picked, sampling from those with visibly different appearances, and cultured in 3 or 5 mL of LB overnight, shaking at 37°C. Cultures were then passaged with a 1:100 dilution into 3 or 5mL of LB and cultured for another 7h, shaking at 37°C. DNA from these cultures was miniprepped using the QIAprep Spin Miniprep kit (Qiagen 27104) and analyzed on a 15% TBE-urea (Thermo Fisher EC6885) gel to visualize possible RT-DNA bands. Based on the length of previously identified retron RT-DNA^22^, bands between 40 and 200 base pairs were considered as a potential signature of a retron.

### Isolating RT-DNA and Sequencing

In samples with potential RT-DNA bands, DNA was isolated from cultures using the Qiagen kit and single stranded DNA was isolated from the miniprep using the ssDNA/RNA clean & concentrator kit (Zymo Research). RT-DNA was prepped for sequencing by taking the resulting material and extending the 3′ end in parallel with dATP, using terminal deoxynucleotidyl transferase (TdT) (NEB). This reaction was carried out in 1× TdT buffer, with 60 units of TdT and 125 M dATP for 60 s at room temperature with the aim of adding ∼25 adenosines before inactivating the TdT at 70°C for 5 min. Next, a polynucleotide anchored primer was used to create a complementary strand to the TdT extended products using 15 units of Klenow Fragment (3′→5′ exo-) (NEB) in 1× NEB2, 1 mM dNTP and 50 nM of primer containing an Illumina adapter sequence, nine thymines (for the pA extended version) or six guanines (for the pC extended version), and a non-thymine (V) or non-guanine (H) anchor. The product was cleaned using a Qiagen PCR cleanup kit and eluted in 10 ul water. Finally, Illumina adapters were ligated on at the 3′ end of the complementary strand using 1× TA Ligase Master Mix (NEB). All products were indexed and sequenced on an Illumina MiSeq instrument.

### Whole Genome Sequencing and Identification of Retron Operons

Genomic DNA was isolated from overnight cultures of host bacteria using the Zymo Quick DNA/RNA Miniprep Kit (Zymo D7001). Genomic DNA was then tagmented with Tn5 transposase preloaded with Illumina sequencing adapters, indexed, and sequenced on a NextSeq 2000. Whole genome contigs for *Pseudomonas peli, Aeromonas salmonicida*, and *Sphingobacter daejeonense* samples were assembled using Geneious Prime software. Sequencing data for the remaining samples included more than 5.5 million reads, so we used a Velvet plugin instead to assemble whole genome contigs more quickly. In the *Huaxiibacter chinensis* and *Serratia quinivorans* samples, sequencing yielded 40 million and 24 million reads, respectively, but only 10% and 50%, respectively, were used for Velvet assemblies.

RT-DNA reads were mapped to whole genome data to locate the potential retron operon. Genomes were additionally analyzed by PADLOC (Prokaryotic Antiviral Defense Locator)^28^, which identifies defense systems, including retrons, based on known motifs. PADLOC also identifies the likely start and end of accessory and RT genes.

In samples where whole genome data was not collected or where assembled contigs did not contain the entire retron operon, genomic DNA was tagmented using Tn5 preloaded with Illumina sequencing adapters and amplified to enrich for retron-containing fragments. To perform this PCR, two primers were designed to bind the known RT-DNA sequence facing opposite directions to capture both ends of the retron. Another two primers were designed to bind the two Illumina adapters at each end of the fragment. Four PCR reactions were then performed, one for every combination of RT-DNA-binding primer and adapter-binding primer. The enriched sample was then sequenced using either MiSeq or Sanger sequencing of the resulting amplicons.

When necessary, this process was repeated to capture the entire retron operon with primers moving successively outward from the known part of the retron.

### Identifying New Retron Subtypes in Sequencing Data

In samples with potential RT-DNA, but where PADLOC did not identify full retron systems, potential retron RT genes were identified using the PADLOC, DefenseFinder, and NCBI’s BLAST programs to find sequences with similarity to known retron RTs. Possible retron accessory genes were identified by studying both their direction and gene functions predicted by BLAST and EMBL-EBI’s Interpro tool.

### Finding homologs of Retron Sda1

To investigate the unusual genomic context of Retron-Sda1, we conducted a comprehensive homology and phylogenetic analysis. Initially, protein sequences corresponding to the reverse transcriptase (RT) and DUF3037 domains of Retron-Sda1 were extracted from the assembled genome contigs. These sequences were used as queries in a BLASTp search against the NCBI non-redundant (nr) protein database [https://ftp.ncbi.nlm.nih.gov/blast/db/FASTA/nr.gz] to identify homologous proteins across diverse bacterial taxa. Multiple sequence alignments of the retrieved RT and DUF3037 protein sequences were performed using MAFFT v7.475 with the G-INS-i algorithm^32^. Phylogenetic trees were constructed using FastTree v2.1^33^ with default parameters. To examine the genomic context of the identified homologs, we utilized the NCBI Entrez Programming Utilities^34^ API to retrieve genomic neighborhoods surrounding each homologous RT and DUF3037 gene. Genomic regions extending ±10 kilobases from each gene were extracted and analyzed for co-localized genes encoding HipA-like kinases and DUF3037 domain-containing proteins.

Defense system annotation was carried out using PADLOC v2.0.0^27^, This analysis was facilitated by custom Python scripts utilizing the Biopython library^35^. Genomic neighorhoods were visualized using a custom JavaScript code and made availale as a portable HTML file.

### Cloning Retron Plasmids and Verifying RT-DNA Production in *E. coli*

Predicted retron operons were amplified by PCR from their host bacterial genomes and cloned into a pET-21(+) backbone (ColE1 ori, kanamycin resistance) in NEB 5-alpha Competent *E. coli*. Plasmids were verified with Sanger sequencing and then transformed into BL21^*ΔEco1*^ (bSLS.114^7^, Addgene #191530). Colonies were picked into 3 mL of LB, grown overnight (shaking at 37°C), passaged at a 1:100 dilution into 3 mL of LB, and then induced with working concentrations of 1 mM IPTG (GoldBio) and 2 mg/ml L-arabinose (GoldBio). After 7 hrs of culture with induction, DNA was miniprepped using the QIAprep Spin Miniprep kit and RT-DNA bands were visualized using electrophoresis on 15% TBE-urea gels (Thermo Fisher EC6885).

### Phage Strains and Propagation

Lambda and T5 phages were initially propagated from ATCC stocks (Lambda WT #23724-B2, T5 #11303-B5). T4 phage was drawn from a previously established stock^30^. MG1655 *E. coli* culture was infected with phage and grown overnight. The culture was then centrifuged for 10 min at 3,434*g* and the supernatant was filtered at 0.2 μm to remove bacterial remnants. Lysate titer was determined using the full plate plaque assay method as described by Kropinski et al^36^. We used a strictly lytic version of lambda carrying two early stop codons in the cI gene, responsible for lysogeny control, to ensure the phage was strictly lytic (lambda ΔcI)^30^. This lambda strain additionally contains a genomic deletion between positions 21738 and 27723 that includes genes involved in lysogeny control. Lambda ΔcI was used for all phage defense experiments. Bas51 phage was propagated from the Leibniz Institute DSMZ stock (https://www.dsmz.de/collection/catalogue/details/culture/DSM-112929), and was originally isolated in the BASEL phage collection^37^.

### Testing Retron Defense Against *E. Coli* Phages

MG1655 cells were transformed with plasmids containing the retron operons and plated. Transformed colonies were picked and cultured overnight in 3 mL of LB (shaking at 37°C) then passaged in 3 mL of MMB (LB medium supplemented with 0.1 mM MnCl2 and 5 mM MgCl2) for 3-5 hrs. 200 µL of culture was plated in 3 mL of MMB soft agar in each well of a 4-well MMB agar plate. Lambda, T4, T5, and Bas51 phages were serially diluted to 10^−4^, 10^−5^, 10^−6^, and 10^−7^ in MMB media. 5 µL of each dilution was spotted on these plates. After the spots have completely dried, plates were incubated at 37°C overnight. Resultant plaques were counted to determine the titer (pfu/mL) of each phage strain on each retron-expressing bacterial lawn.

### Isolating Mutant Phages and Identifying Retron Trigger Genes

For retron and phage combinations where defense occurred, phages and retrons were plated on MMB soft agar at the lowest phage dilution where few or no plaques were visible. Plaques were then picked from plates and, along with the wild type phage, propagated in a culture of the MG1655 transformed with the given retron. These cultures were plated in MMB soft agar and grown overnight. SM buffer (100 mM NaCl, 8 mM MgSO4, 50 mM Tris-HCl pH 7.5) was added to submerge the wells and left for 4 hrs at 4°C to extract phages. The buffer was then removed and filtered with a 0.2 μm filter. For amplification-free sequencing, extracellular DNA was removed through DNase I treatment, with 20 U of DNase I (NEB, M0303S) per 1 mL of phage lysate, incubated at room temperature for 15 min and then inactivated at 75°C for 5 min. Phage were then lysed and DNA extracted using the Norgen Phage DNA Isolation Kit (Norgen, 46800). Phage mutant samples were prepped for nanopore sequencing using the Ligation Sequencing Kit (SQK-LSK109) and Native Barcoding Expansion (EXP-NBD196) from Oxford Nanopore Technologies (ONT). Nanopore sequencing was performed using an R9.4.1 flow cell (FLO-MIN106D) on a MinION device. Wild type phage samples were prepped using the Ligation Sequencing Kit (SQK-LSK114) and sequenced using an R10.4.1 flow cell (FLO-PRO114M) on a PromethION device. Base calling was performed using ONT’s Guppy (for phage mutants) or Dorado (for wild type phage) Basecaller software on the high accuracy setting.

Nanopore sequencing reads for mutant and wild type phage were assembled independently using the Geneious Prime software and consensus sequences were mapped to reference genomes to identify mutations. Genes containing mutations in multiple phage escapees for a given retron were considered potential trigger genes. To confirm potential trigger genes, wild type phage was engineered to incorporate the individual mutations seen in the escapee (see *Phage Recombineering* below). These engineered phage mutants were then tested for defense in the same retron-expressing bacterial strain they were isolated from (see *Testing Retron Defense Against* E. coli *Phages* above).

### Phage Recombineering

To make each phage mutant, a recombitron plasmid (as described in Fishman et al.^30^) with a 90 nt editing donor was co-transformed with pORTMAGE (contains CspRecT and mutL E32K accessory genes for recombineering) into a modified strain of *E. coli* MG1655 with inactivated *exoI* and *recJ* genes, as well as an arabinose-inducible T7 polymerase (bMS.346^7^, Addgene #220588). Individual transformants were picked from plates and grown in 3 mL LB media overnight in a shaking incubator at 37°C. The following day, these bacterial editor cultures were diluted 1:100 in 3 mL MMB media containing 1 mM IPTG (GoldBio), 2 mg/ml L-arabinose (GoldBio), and 1 mM m-toluic acid (Sigma-Aldrich) to express the recombineering machinery.

Passaged cultures were grown for an additional 2-3 hrs at 37°C, then diluted to an OD600 of 0.2 in a fresh culture of 3 mL MMB with inducers. Wild type phage lysate was spiked into these cultures at a MOI of 0.1. The infected cultures were then grown for 16-18 hrs in a shaking incubator at 37°C. Cultures were then centrifuged for 10 min at 3,434*g* and the supernatant was filtered at a 0.2 μm pore size to obtain clarified phage lysate. To increase the percent of edited phage in the population, this recombineering protocol was repeated using the newly isolated phage lysate as the spike-in for a fresh batch of bacterial editors. Editing rates were monitored after every round of recombineering by PCR amplifying the phage lysate with primers flanking the edit site and then Sanger sequencing those amplicons.

After two rounds of recombineering, individual phage mutants were isolated by plaquing on a lawn of bacterial editors. To do this, editors were grown overnight and passaged in 3 mL MMB with inducers for 3-5 hrs. Edited phage lysate was diluted 1:10000 in MMB media. Then, 5 µL of diluted lysate was spiked into 1 mL of passaged editor culture, all of which was mixed into 20 mL of soft agar and poured onto a LB plate. Soft agar plates were incubated at 37°C overnight. The next morning, eight individual plaques were picked for each phage mutant, resuspended in 100 µL of SM Buffer, and left at 4°C for at least 4 hours to allow the phage to disseminate into the buffer. Amplification of the edit site followed by Sanger sequencing was again performed to screen for lysates containing the desired edit. Those lysates were further propagated to obtain high titer phage mutants for use in subsequent experiments.

### Engineering Retrons for Recombineering in *E. coli*

To edit bacterial genomes using recombineering, retrons were modified to encode a recombineering donor in the reverse transcribed region of the ncRNA. This 70 nt donor incorporates a 10 nt deletion in the *lacZ* gene. Effector proteins were excluded from retrons for recombineering. The modified retrons were cloned into the pACYC-Duet1 backbone (p15A ori, chloramphenicol resistance) where they are co-expressed with CspRecT under the constitutive J23115 promoter. These plasmids were then transformed into bMS.346 cells and immediately inoculated into 5 mL LB cultures, which were grown overnight, shaking at 37°C. The next day, 25 µl of culture was collected, mixed with 25 µl of water, and boiled at 95°C for 10 min. The same cultures were then passaged at a 1:100 dilution in fresh 5 mL LB and grown overnight again, after which another set of boiled lysates was collected. 0.5 µl of each boiled lysate from Day 1 and Day 2 of editing was then used as a template in 25 µl PCR reactions with primers flanking the edit site, which additionally contained adapters for Illumina sequencing preparation. These amplicons were indexed and sequenced on an Illumina MiSeq or NextSeq 2000 instrument. Sequencing reads were processed with CRISPResso2^38^ (https://github.com/pinellolab/CRISPResso2) to quantify the percentage of precisely edited genomes. The following equation was used to calculate percent editing:

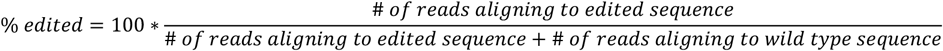

## ACKNOWLEDGEMENTS

Work was supported by funding from the National Science Foundation (MCB 2137692), the Gary and Eileen Morgenthaler Fund, the Gordon and Betty Moore Foundation, and the Robert J Kleberg, Jr. and Helen C. Kleberg Foundation. S.L.S. is a Chan Zuckerberg Biohub – San Francisco Investigator. K.Z. was supported by a National Science Foundation Graduate Research Fellowship and a UCSF Discovery Fellowship.

## AUTHOR CONTRIBUTIONS

Author contributions follow the CRedIT taxonomy (https://www.elsevier.com/researcher/author/policies-and-guidelines/credit-author-statement)

K.N.: Methodology, Investigation, Validation, Writing.

K.Z.: Methodology, Investigation, Validation, Writing, Supervision. M.R.M.: Formal Analysis, Software, Writing

M.R-M.: Methodology, Supervision.

S.L.S.: Conceptualization, Methodology, Formal Analysis, Writing, Visualization, Supervision, Project Administration, Funding Acquisition.

### DATA AND CODE AVAILABILITY

Sequencing data associated with this study are available in the NCBI SRA (PRJNA1196507) https://www.ncbi.nlm.nih.gov/sra/PRJNA1196507

***Supplementary File 1 (uploaded separately):*** Interactive HTML file showing the genomic neighborhoods of all DUF3037 homologs.

***Supplementary File 2 (uploaded separately):*** Interactive HTML file showing the genomic neighborhoods of the six Sda1-like retrons.

## SUPPLEMENTARY MATERIALS

**Supplementary Figure 1.**
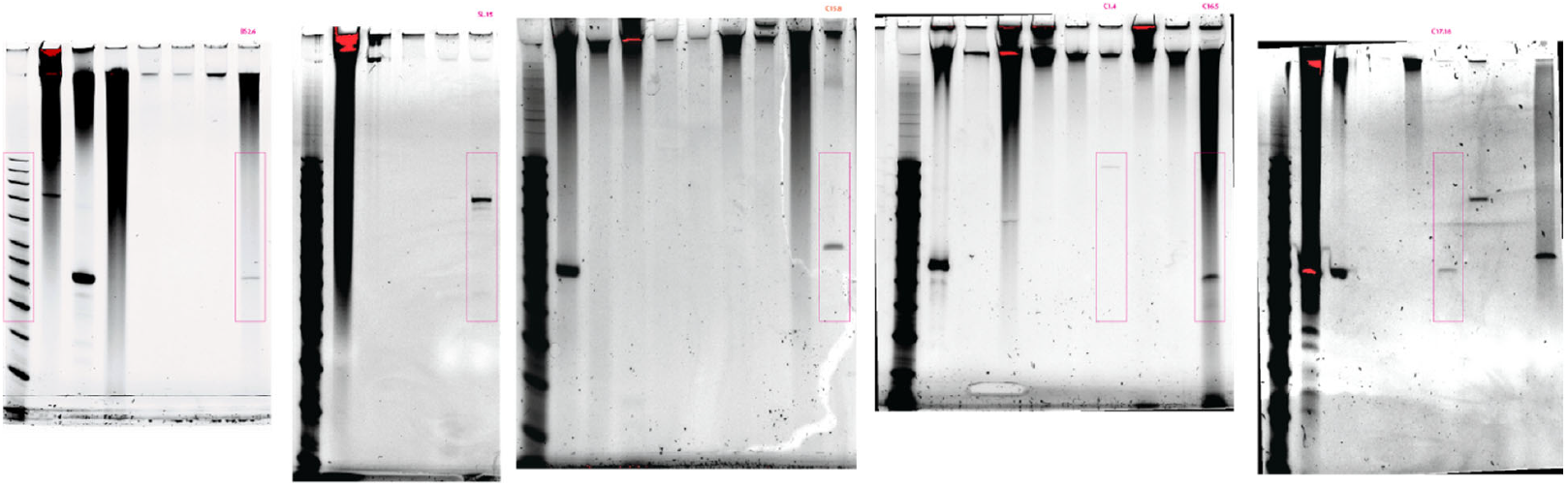
Related to Figure 1c. *Uncropped gels.*

**Supplementary Figure 2.**
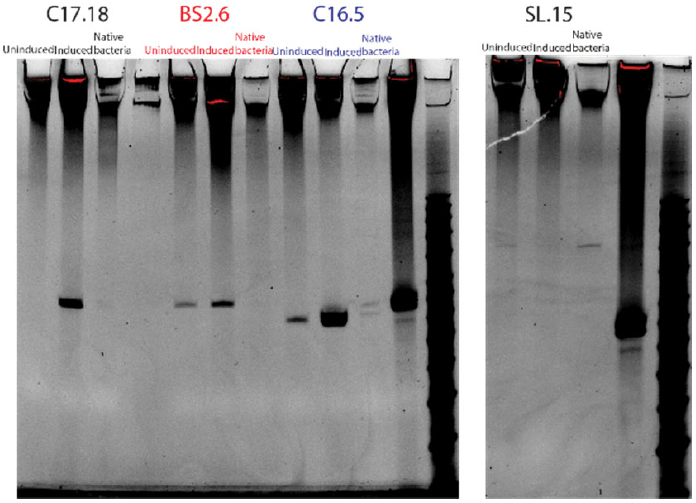
Related to Figure 2b. *Uncropped gels.*

**Supplementary Table 1.** Sample characteristics and retrons identified.

| Soil |  |  |
| --- | --- | --- |
| Code | Characteristics | Retron |
| GSS1 | Small garden in urban area |  |
| GSS2 | Small garden in urban area |  |
| Koret Quad | Park in an urban area |  |
| BS1 | Surface level soil from a park |  |
| BS2 | Soil 1.5 ft below ground from a park | Retron-Squ1 |
| CS1 | High altitude, dry soil |  |
| CS2 | High altitude, dry soil |  |
| CS3 | High altitude, dry soil |  |
| C1 | Grassland | Retron-Ppe1 |
| C4 | High altitude forest |  |
| C6 | High altitude forest |  |
| C10 | High altitude glacial lake |  |
| C11 | High altitude, mountainous terrain |  |
| C12 | High altitude, mountainous terrain |  |
| C13 | High altitude, mountainous terrain |  |
| C17 | High altitude, mountainous terrain | Retron-Hch1 |
| C19 | High altitude glacial lake |  |
| VI ab | Urban area |  |
| VI 2a | Urban area |  |
| Water |  |  |
| Code | Characteristics | Retron |
| SL | Freshwater pond | Retron-Bpu1 |
| FT | Aquarium with fish |  |
| FC | Saltwater lagoon |  |
| C2 | High altitude freshwater stream |  |
| C3 | High altitude glacial lake |  |
| C5 | High altitude glacial lake |  |
| C7 | High altitude glacial lake |  |
| C8 | High altitude glacial lake |  |
| C9 | High altitude glacial lake |  |
| C14 | High altitude glacial lake |  |
| C16 | High altitude glacial lake | Retron-Asa1 |
| C15 | High altitude glacial lake | Retron-Sda1 |
| C18 | High altitude freshwater stream |  |

**Supplementary Table 2.** Additional Statistical Details.

| Figure 3 |  |  |  |  |  |
| --- | --- | --- | --- | --- | --- |
| Panel | Biological Replicates | Comparison | Test | P value | P value summary |
| 4a | 3 | Effect of retron | Kruskal-Wallis test | 0.3385 | ns |
| 4c | 3 | Effect of retron | Kruskal-Wallis test | 0.0963 | ns |
| 4e | 3 | Effect of retron | Kruskal-Wallis test | 0.9084 | ns |
| 4g | 3-6 | Effect of retron | Kruskal-Wallis test | 0.0047 | ** |
|  |  | follow-up: no retron vs Asa1 | Dunn's multiple comparisons test (corrected) | 0.0631 | ns |
|  |  | follow-up: no retron vs Squ1 | Dunn's multiple comparisons test (corrected) | 0.0009 | *** |
|  |  | follow-up: no retron vs Exx1 | Dunn's multiple comparisons test (corrected) | >0.9999 | ns |
|  |  | follow-up: no retron vs Bpu1 | Dunn's multiple comparisons test (corrected) | 0.3338 | ns |
| Figure 4 |  |  |  |  |  |
| Panel | Biological Replicates | Comparison | Test | P value | P value summary |
| 5c | 3 | Effect of retron | Two-way ANOVA | <0.0001 | **** |
|  |  | Effect of phage | Two-way ANOVA | 0.1244 | ns |
|  |  | Effect of interaction | Two-way ANOVA | <0.0001 | **** |
|  |  | no retron: wt vs. methyl D69G Bas51 | Tukey's multiple comparisons test (corrected) | 0.0006 | *** |
|  |  | no retron: wt vs. methyl D79G Bas51 | Tukey's multiple comparisons test (corrected) | 0.0011 | ** |
|  |  | no retron: wt vs. methyl T83P Bas51 | Tukey's multiple comparisons test (corrected) | <0.0001 | **** |
|  |  | no retron: wt vs. glyco R87* Bas51 | Tukey's multiple comparisons test (corrected) | 0.0046 | ** |
|  |  | wt Bas51: no retron vs. Asa1 | Tukey's multiple comparisons test (corrected) | <0.0001 | **** |
|  |  | wt Bas51: no retron vs. Squ1 | Tukey's multiple comparisons test (corrected) | <0.0001 | **** |
|  |  | methyl D69G: no retron vs. Asa1 | Tukey's multiple comparisons test (corrected) | >0.9999 | ns |
|  |  | methyl D69G: no retron vs. Squ1 | Tukey's multiple comparisons test (corrected) | 0.7463 | ns |
|  |  | methyl D79G: no retron vs. Asa1 | Tukey's multiple comparisons test (corrected) | >0.9999 | ns |
|  |  | methyl D79G: no retron vs. Squ1 | Tukey's multiple comparisons test (corrected) | 0.9472 | ns |
|  |  | methyl T83P: no retron vs. Asa1 | Tukey's multiple comparisons test (corrected) | 0.9472 | ns |
|  |  | methyl T83P: no retron vs. Squ1 | Tukey's multiple comparisons test (corrected) | 0.7463 | ns |
|  |  | glyco R87*: no retron vs. Asa1 | Tukey's multiple comparisons test (corrected) | 0.491 | ns |
|  |  | glyco R87*: no retron vs. Squ1 | Tukey's multiple comparisons test (corrected) | 0.0001 | *** |

**Supplementary Table 3.**
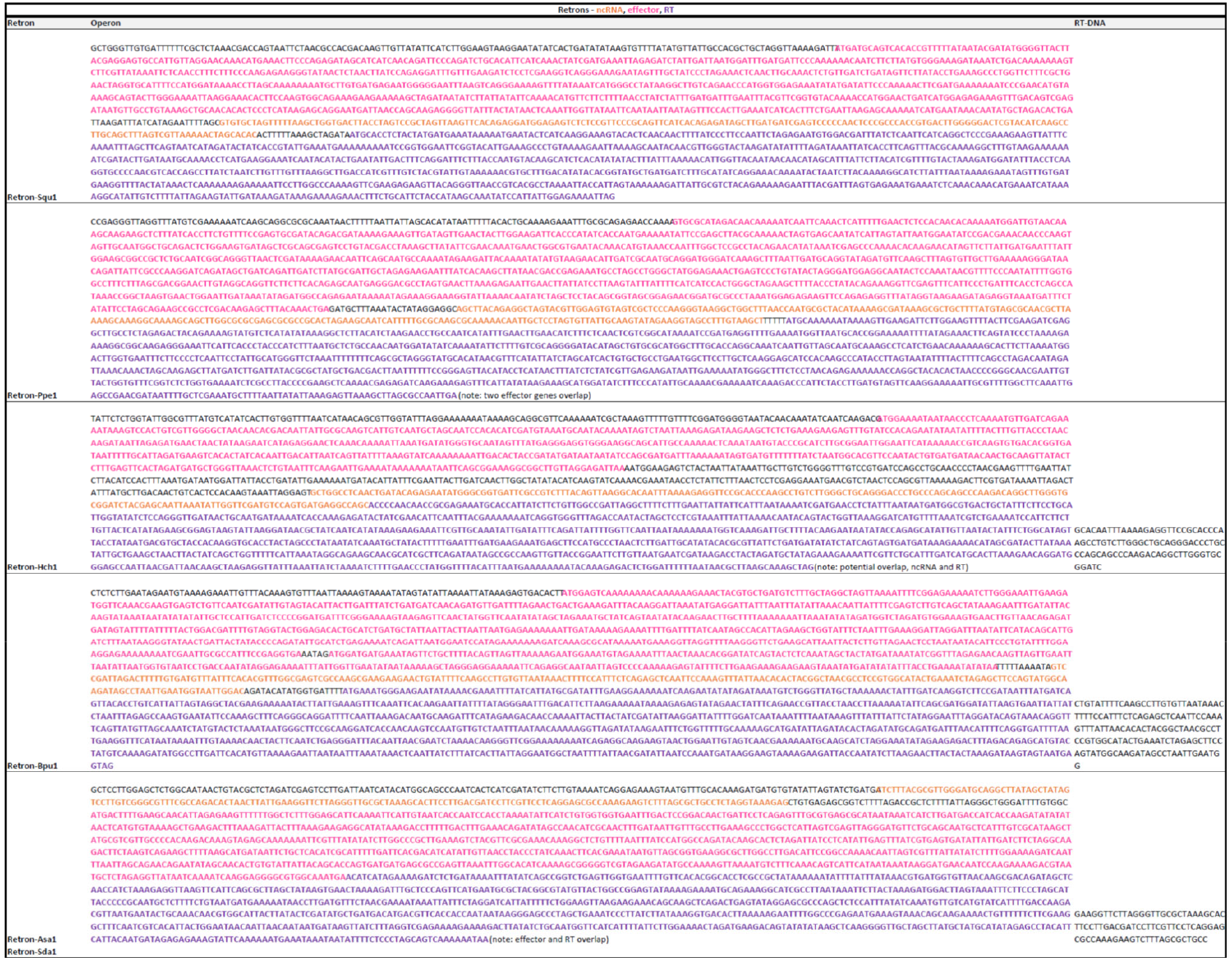
Retron Operons.

## REFERENCES

1 Bobonis, J. et al. Bacterial retrons encode phage-defending tripartite toxin-antitoxin systems. Nature 609, 144–150, doi:10.1038/s41586-022-05091-4 (2022).

2 Millman, A. et al. Bacterial Retrons Function In Anti-Phage Defense. Cell 183, 1551-1561.e1512, doi:10.1016/j.cell.2020.09.065 (2020).

3 Gao, L. et al. Diverse enzymatic activities mediate antiviral immunity in prokaryotes. Science 369, 1077–1084, doi:10.1126/science.aba0372 (2020).

4 Bhattarai-Kline, S. et al. Recording gene expression order in DNA by CRISPR addition of retron barcodes. Nature 608, 217–225, doi:10.1038/s41586-022-04994-6 (2022).

5 Vibhute, M. A. et al. Intracellular Expression of a Fluorogenic DNA Aptamer Using Retron Eco2. bioRxiv, 2024.2005.2021.595248, doi:10.1101/2024.05.21.595248 (2024).

6 Liu, J. et al. Generation of DNAzyme in Bacterial Cells by a Bacterial Retron System. ACS Synth Biol 13, 300–309, doi:10.1021/acssynbio.3c00509 (2024).

7 Lopez, S. C., Crawford, K. D., Lear, S. K., Bhattarai-Kline, S. & Shipman, S. L. Precise genome editing across kingdoms of life using retron-derived DNA. Nat Chem Biol 18, 199–206, doi:10.1038/s41589-021-00927-y (2022).

8 Farzadfard, F. & Lu, T. K. Genomically encoded analog memory with precise in vivo DNA writing in living cell populations. Science 346, 1256272–1256272, doi:10.1126/science.1256272 (2014).

9 Schubert, M. G. et al. High-throughput functional variant screens via in vivo production of single-stranded DNA. PNAS 118, e2018181118, doi:10.1073/pnas.2018181118 (2021).

10 Kong, X. et al. Precise genome editing without exogenous donor DNA via retron editing system in human cells. Protein & cell 12, 899–902, doi:10.1007/s13238-021-00862-7 (2021).

11 Zhao, B., Chen, S. A., Lee, J. & Fraser, H. B. Bacterial Retrons Enable Precise Gene Editing in Human Cells. The CRISPR journal 5, 31–39, doi:10.1089/crispr.2021.0065 (2022).

12 Cattle, M. A. et al. An enhanced Eco1 retron editor enables precision genome engineering in human cells from a single-copy integrated lentivirus. bioRxiv, 2024.2008.2005.606586, doi:10.1101/2024.08.05.606586 (2024).

13 Buffington, J. D. et al. Discovery and Engineering of Retrons for Precise Genome Editing. bioRxiv, 2024.2007.2021.604473, doi:10.1101/2024.07.21.604473 (2024).

14 Lim, H. et al. Multiplex Generation, Tracking, and Functional Screening of Substitution Mutants Using a CRISPR/Retron System. ACS Synth Biol 9, 1003–1009, doi:10.1021/acssynbio.0c00002 (2020).

15 Liu, W. et al. Retron-mediated multiplex genome editing and continuous evolution in Escherichia coli. Nucleic Acids Res 51, 8293–8307, doi:10.1093/nar/gkad607 (2023).

16 Ni, Y. et al. Reducing competition between <em>msd</em> and genomic DNA significantly improved the editing efficiency of the retron editing system. bioRxiv, 2024.2006.2004.597346, doi:10.1101/2024.06.04.597346 (2024).

17 Jiang, W. et al. High-efficiency retron-mediated single-stranded DNA production in plants. Synthetic biology (Oxford, England) 7, ysac025, doi:10.1093/synbio/ysac025 (2022).

18 Crawford, K. D., Khan, A. G., Lopez, S. C., Goodarzi, H. & Shipman, S. L. High throughput variant libraries and machine learning yield design rules for retron gene editors. Nucleic Acids Res, doi:10.1093/nar/gkae1199 (2024).

19 Yee, T., Furuichi, T., Inouye, S. & Inouye, M. Multicopy single-stranded DNA isolated from a gram-negative bacterium, Myxococcus xanthus. Cell 38, 203–209, doi:10.1016/0092-8674(84)90541-5 (1984).

20 Dhundale, A., Lampson, B., Furuichi, T., Inouye, M. & Inouye, S. Structure of msDNA from Myxococcus xanthus: evidence for a long, self-annealing RNA precursor for the covalently linked, branched RNA. Cell 51, 1105–1112, doi:10.1016/0092-8674(87)90596-4 (1987).

21 Mestre, M. R., González-Delgado, A., Gutiérrez-Rus, L. I., Martínez-Abarca, F. & Toro, N. Systematic prediction of genes functionally associated with bacterial retrons and classification of the encoded tripartite systems. Nucleic Acids Res 48, 12632–12647, doi:10.1093/nar/gkaa1149 (2020).

22 Khan, A. G. et al. An experimental census of retrons for DNA production and genome editing. Nature biotechnology, doi:10.1038/s41587-024-02384-z (2024).

23 Palka, C., Fishman, C. B., Bhattarai-Kline, S., Myers, S. A. & Shipman, S. L. Retron reverse transcriptase termination and phage defense are dependent on host RNase H1. Nucleic Acids Res 50, 3490–3504, doi:10.1093/nar/gkac177 (2022).

24 Wang, Y. et al. Cryo-EM structures of Escherichia coli Ec86 retron complexes reveal architecture and defence mechanism. Nature microbiology 7, 1480–1489, doi:10.1038/s41564-022-01197-7 (2022).

25 Carabias, A. et al. Retron-Eco1 assembles NAD(+)-hydrolyzing filaments that provide immunity against bacteriophages. Molecular cell 84, 2185-2202.e2112, doi:10.1016/j.molcel.2024.05.001 (2024).

26 Stokar-Avihail, A. et al. Discovery of phage determinants that confer sensitivity to bacterial immune systems. Cell 186, 1863-1876.e1816, doi:10.1016/j.cell.2023.02.029 (2023).

27 Caigoy, J. C. et al. Genetic Characterization of a Novel Retron Element Isolated from Vibrio mimicus. Microbiology and immunology 69, 1–9, doi:10.1111/1348-0421.13181 (2025).

28 Payne, L. J. et al. PADLOC: a web server for the identification of antiviral defence systems in microbial genomes. Nucleic Acids Res 50, W541–w550, doi:10.1093/nar/gkac400 (2022).

29 DeWeirdt, P. C., Mahoney, E. M. & Laub, M. T. DefensePredictor: A Machine Learning Model to Discover Novel Prokaryotic Immune Systems. bioRxiv, 2025.2001.2008.631726, doi:10.1101/2025.01.08.631726 (2025).

30 Fishman, C. B. et al. Continuous multiplexed phage genome editing using recombitrons. Nature biotechnology, doi:10.1038/s41587-024-02370-5 (2024).

31 González-Delgado, A., Lopez, S. C., Rojas-Montero, M., Fishman, C. B. & Shipman, S. L. Simultaneous multi-site editing of individual genomes using retron arrays. Nat Chem Biol, doi:10.1038/s41589-024-01665-7 (2024).

32 Katoh, K. & Standley, D. M. MAFFT multiple sequence alignment software version 7: improvements in performance and usability. Molecular biology and evolution 30, 772–780, doi:10.1093/molbev/mst010 (2013).

33 Price, M. N., Dehal, P. S. & Arkin, A. P. FastTree 2--approximately maximum-likelihood trees for large alignments. PLOS ONE 5, e9490, doi:10.1371/journal.pone.0009490 (2010).

34 (US), N. C. f. B. I. <https://www.ncbi.nlm.nih.gov/books/NBK25501/> (2010-).

35 Cock, P. J. et al. Biopython: freely available Python tools for computational molecular biology and bioinformatics. Bioinformatics (Oxford, England) 25, 1422–1423, doi:10.1093/bioinformatics/btp163 (2009).

36 Kropinski, A. M., Mazzocco, A., Waddell, T. E., Lingohr, E. & Johnson, R. P. Enumeration of bacteriophages by double agar overlay plaque assay. Methods in molecular biology (Clifton, N.J.) 501, 69–76, doi:10.1007/978-1-60327-164-6_7 (2009).

37 Maffei, E. et al. Systematic exploration of Escherichia coli phage-host interactions with the BASEL phage collection. PLoS biology 19, e3001424, doi:10.1371/journal.pbio.3001424 (2021).

38 Clement, K. et al. CRISPResso2 provides accurate and rapid genome editing sequence analysis. Nature biotechnology 37, 224–226, doi:10.1038/s41587-019-0032-3 (2019).

